# The Relation between Alpha/Beta Oscillations and the Encoding of Sentence induced Contextual Information

**DOI:** 10.1101/501437

**Authors:** René Terporten, Jan-Mathijs Schoffelen, Bohan Dai, Peter Hagoort, Anne Kösem

## Abstract

Within the sensory domain, alpha/beta oscillations have been frequently linked to the prediction of upcoming sensory input. Here, we investigated whether oscillations at these frequency bands serve as a neural marker in the context of linguistic input prediction as well. Specifically, we hypothesized that if alpha/beta oscillations do index language prediction, their power should modulate during sentence processing, indicating stronger engagement of underlying neuronal populations involved in the linguistic prediction process. Importantly, the modulation should monotonically relate to the degrees of predictability of incoming words based on past context. Specifically, we expected that the more predictable the last word of a sentence, the stronger the alpha/beta power modulation. To test this, we measured neural responses with magnetoencephalography of healthy individuals (of either sex) during exposure to a set of linguistically matched sentences featuring three distinct levels of sentence context constraint (high, medium and low constraint). We observed fluctuations in alpha/beta power before last word onset, and also modulations in M400 amplitude after last word onset that are known to gradually relate to semantic predictability. In line with previous findings, the M400 amplitude was monotonically related to the degree of context constraint, with a high constraining context resulting in the strongest amplitude decrease. In contrast, alpha/beta power was non-monotonically related to context constraints. The strongest power decrease was observed for intermediate constraints, followed by high and low constraints. While the monotonous M400 amplitude modulation fits within a framework of prediction, the non-monotonous oscillatory results are not easily reconciled with this idea.

**SIGNIFICANCE STATEMENT:** Neural activity in the alpha (8-10Hz) and beta (16-20) frequency ranges have been related to the prediction of upcoming sensory input. It remains still debated whether these frequency bands relate to language prediction as well. In this magnetoencephalography study, we recorded alpha/beta oscillatory activity while participants listened to sentences whose ending had varying degree of predictability based on past linguistic information. Our results show that alpha/beta power modulations were non-monotonically related to the degree of linguistic predictability: the strongest modulation of alpha/beta power was observed for intermediate levels of linguistic predictability during sentence reading. Together, the results emphasize that alpha/beta oscillations cannot directly be linked to predictability in language, but potentially relate to attention or control operations during language processing.

## INTRODUCTION

Sentence level language comprehension results from dynamic cognitive processes which combine and unify smaller linguistic units to create meaning (Cairns et al., 1981; Glucksberg et al., 1986; Hagoort, 2016; Morris, 1994; Moss & Marslen-Wilson, 1993; Rommers et al., 2013). These cognitive processes occur online, while the sentence unfolds, instantiating unified meaning which relates to the computation of semantics, spanning the whole utterance. During this process, a context representation is compared and updated on a moment to moment basis. The bias provided by the momentarily established context alters subsequent linguistic processing (Federmeier, 2007; Frank & Willems, 2017; Xu et al., 2005) and is classically marked on a neuronal level by the N400 component (M400 in MEG studies) (Janssen et al., 2017; Kutas & Federmeier, 2011; Tromp et al., 2017). A monotonic decrease of the N400 amplitude has been associated with increasing context constraints (DeLong et al., 2005; Diaz & Swaab, 2007; Freunberger & Roehm, 2017; Ito et al., 2016; Kutas & Federmeier, 2011; Van Petten & Luka, 2012). While the N400 response is observed after target word onset, its presence is compatible with the possibility that sentence context constraints alter predictions that are encoded *prior* to target word occurrence.

Prediction in its minimal sense can be understood as changes in brain *states* in response to contextual information which facilitate the processing of new input (Kuperberg & Jaeger, 2015). Recent evidence suggests that neural rhythmic activity in the alpha (8-12 Hz) and low-beta (16-20 Hz) ranges could be involved in the prediction of linguistic input during sentence processing. Alpha/beta oscillations are hypothesized to play a crucial role for the prediction of upcoming sensory input (Arnal & Giraud, 2012; Lewis et al., 2016), and to constitute top-down mechanisms that shape the communication of sensory information between distant neural networks (Bastos et al., 2015; Bonnefond et al., 2017; Fries, 2015). Beyond sensory processing, recent theory and evidence suggests that alpha/beta oscillations would also be involved in linguistic prediction (Kuperberg & Jaeger, 2015; Lewis & Bastiaansen, 2015; Lewis et al., 2015). Decreases in alpha power (Lam et al., 2016; Piai et al., 2017; Rommers et al., 2017; Wang et al., 2017) and low-beta power (Bastiaansen & Hagoort, 2015; Piai et al., 2017; Wang et al., 2017) have previously been linked to the processing of sentential context constraints. Specifically, the power decrease has been found to be stronger when sentential context is highly predictive of the last word of the sentence (i.e. when cloze probability is high) than when the prediction of the last word cannot be made based on past context (i.e. when cloze probability is low) (Bastiaansen & Hagoort, 2015; Lam et al., 2016; Rommers et al., 2017; Piai et al., 2017; Wang et al., 2017; Wang et al., 2012; Weiss & Mueller, 2012; Willems et al., 2008). Yet, the evidence for alpha/beta oscillations in language prediction is still debated, as it has only been observed for strong cloze probability situations (Piai et al., 2017; Rommers et al., 2017; Wang et al., 2017). If alpha/beta power reflect the degree of predictability of an upcoming word, we hypothesized that alpha/beta power should gradually decrease with higher cloze probability. A non-monotonic decrease of alpha/beta power would fit with alternative interpretations of the findings, such that alpha/beta oscillations rather reflect memory operations (Piai et al., 2015) or attentional control (Jensen & Mazaheri, 2010) during sentence processing.

To test this, we introduced sentences with varying degrees of context constraints. Participants passively read sentences belonging to either a low (LC), medium (MC) or high (HC) context constraining condition (Table 1). Neuronal activity was measured online using magnetoencephalography (MEG), before and after display of a target word. We predicted that pre-stimulus alpha and beta power would differ between different conditions of predictability. The power decrease is expected to be strongest for the HC, followed by the MC and LC condition (Fig. 1).

**Figure 1.**
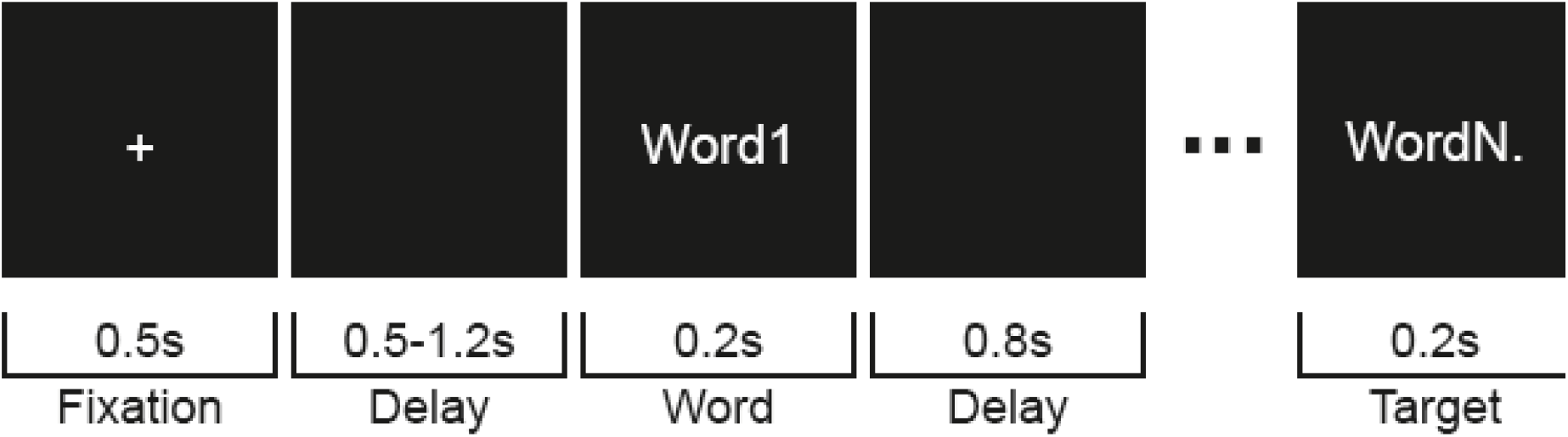
A schematic display of a trial procedure. A trial began with the display of a fixation period, followed by a blank screen. Subsequently the sentence was visually displayed by a word by word presentation, up to the final word as indexed by the period. Between words, a black screen served as delay before a subsequent word was shown.

**Table 1.** Example Dutch sentence triplet from the final stimulus set with its English translation. The context constraining conditions were manipulated by changing one context constraining word.

| Condition | Stimulus |
| --- | --- |
| HC | (NL) Op dit *gebouw* heb je een goed uitzicht.<br>(EN) On this *building* you got a good view. |
| MC | (NL) Op deze *toren* heb je een goed uitzicht.<br>(EN) On this *tower* you got a good view. |
| LC | (NL) In deze *wijk* heb je een goed uitzicht.<br>(EN) In this *area* you got a good view. |

## MATERIALS & METHODS

### Participants

In total, thirty-five students (mean age 24 years, range 18 - 43; 16 males) took part in the experiment. All participants provided their informed consent in accordance with the declaration of Helsinki, and the local ethics committee (CMO region Arnhem-Nijmegen). The participants were all Dutch native speakers, right-handed, had normal or corrected-to-normal vision and none of them suffered from neurological impairment or dyslexia. Two participants were excluded due to poor behavioral performance. Therefore, thirty-three participants were included for further analyses (mean age 24 years, range 18 - 43; 15 males).

### Stimulus material

The stimulus set consisted of 253 sentence triplets, including 203 critical and 50 filler sentence triplets. Each sentence within a critical triplet belonged to either a high context (HC), medium context (MC), or low context (LC) constraining condition. The different degree of constraint within a triplet was achieved by manipulating only one word, the *context constraining word*, which was always at the same position within a sentence with regard to a triplet (table 1). Across the conditions, these context constraining words were matched with regard to word length (F(2, 606) = 0.784, p = .457, with a Mean (SE) of HC: 7.12 (2.26); MC: 7.1 (2.54); LC: 7.37 (2.61)) and word frequency (F(2, 584) = 1.984, p = .138, with Mean (SE) of HC: 2.4 (0.78); MC: 2.56 (0.87); LC: 2.5 (0.84); based on the Dutch SUBTLEX-NL database (Keuleers, et al., 2010). The stimuli were pretested in a sentence completion task in order to verify the step-like degree of context constraints within a triplet (from high, to medium, to low). For this task - independent from the MEG experiment - a sample of participants (N = 51) were required to complete a sentence presented on a computer screen, for which the final word was missing. Participants performed the experiment with one of three counterbalanced lists. Each list included the same amount of critical sentences from either of the three context constraining conditions. The validation of the conditions was performed in two subsequent steps: first, the degree of context constraint per sentence was evaluated by calculating the percentage of participants that would finish a sentence with the same word. As expected, HC sentences resulted in the highest percentage of participants proposing the same word as cloze (Mean (SE) = 77% (17.74)), followed by MC (Mean (SE) = 50% (18.67)) and LC (Mean (SE) = 28% (11.97)). The three conditions differed significantly from each other with regard to their degree of context constraints (F(2, 606) = 442.842, p < .001).

Second, and in order to create the final stimulus set, the final word from the HC sentences with the highest percentage was chosen as sentence ending for all sentences within a triplet. This approach resulted in cloze probabilities for the final word - the *target word* - that were different from the percentages of the context constraints for the MC and LC conditions. Still, the cloze probabilities differed significantly between conditions (F(2, 606) = 468.155, p < .001), with HC showing the highest cloze probability (Mean (SE) = 77% (17.74)), followed by MC (Mean (SE) = 42% (25.94)) and LC (Mean (SE) = 15% (15.82)).

In the MEG experiment, participants were presented with one of the counterbalanced lists, with an additional set of 50 filler sentences. The filler sentences did not differ between lists but followed a different sentence structure as compared to the critical sentences.

### Experimental procedure

Participants were comfortably seated in a dimly illuminated and magnetically shielded room. All participants were instructed to place their arms on the arm rest of the chair, with access to a button box with their right hand. In front of each participant, at a distance of 80 cm, a screen was located on which all stimulus material was displayed. The words were shown in black, on a grey background. Participants were instructed to silently read the displayed sentences on the screen, and to focus on the content of each sentence. Furthermore, it was highlighted that sometimes a question would be asked about the content of the previous displayed sentence. The participants were then required to answer this question with ‘yes’ or ‘no’ by button press. The answer possibilities (‘yes’/’no’) were randomly displayed on the left or right side of the screen and a left or right button had to be pressed accordingly. These question trials were catch trials, intended to ensure that participants were actively processing the meaning of the sentences, without focusing their attention on the contextual constraints. A trial started with the display of a fixation cross in the middle of the screen for 500 ms. The fixation cross was followed by a blank screen for a random interval of 500-1200 ms. Subsequently, the word-by-word presentation of the sentence began, with each word being displayed for 200 ms, followed by a blank screen of 800 ms. An ISI of 1000 ms was chosen in order to record pre-stimulus alpha and beta activity that is not influenced by the evoked response to the previous displayed word. After a sentence ended, another blank screen occurred for 2000 ms (fig. 1). After that, either a catch question was displayed, with the whole question in the middle of the screen and the yes-no answers randomly split to the left or right side, or the next trial began. In total, participants read 253 sentences (253 trials) in random order, which came from one of three lists, counterbalanced on the three levels of context constraints. The total amount of trials was divided into four blocks, separated by small breaks in-between. The length of a break was self-determined by the participants and the task could be continued by button press. In total, the data acquisition lasted 60 min.

### Data acquisition

All data were acquired using a 275 axial gradiometers CTF Omega MEG system. Horizontal and vertical bipolar EOG as well as ECG were recorded in order to discard eye blinks, eye movements and heart beats contaminated trials. All electrophysiological signals were low-pass filtered at 300 Hz, digitized at 1200 Hz, and stored for off-line analysis. Three coils were placed on the nasion and the left and right ear canal to register the head position with respect to the gradiometers. The coils enabled real-time monitoring of the head position throughout the experiment (Stolk et al., 2013). Next to the MEG recordings, magnetic resonance images (MRIs) were obtained from 32 of the participants with a 1.5T or 3.0T Siemens system. By means of attached markers at the same anatomical locations as the head coils, the MRIs could be aligned to the MEG coordinate system.

### Data preprocessing

All data were analyzed using the open-source Matlab toolbox Fieldtrip (Oostenveld et al., 2011). From the MEG data, a time-window of interest was segmented 2 s before and after the onset of a sentence’s final word for each trial. This segmentation therefore included the blank delay period just before onset of the target word, where the effect of context constraints is expected to occur, and the period after onset of the target word. The segmented data were low-pass filtered at 150 Hz. The 50 Hz line noise components were removed by using a notch filter. Artifact identification and rejection was done in three steps. First, MEG jump and muscle artifacts were identified by visual inspection of amplitude variance over trials. Second, artifacts related to eye-movements and cardiac activity were identified and removed by means of an independent component analysis (fastICA; Hyvärinen & Oja, 2000), followed by backprojection. The independent components were visually inspected and removed from the sensor data, if they resembled heartbeat, eye-movements or blinks (as compared to the recorded EOG and ECG). Third, the resulting data were again visually inspected to remove any remaining artifacts. From this procedure, on average 11% of trials and 1.5% of MEG sensors were excluded from further analysis.

### Event-Related Field (ERF) analysis

Event-related fields were investigated to observe M400 modulations after the last word onset. This correlate is the magnetic counterpart of the classical N400 measured by electroencephalography, and inhabits the same time-course and response properties (Halgren et al., 2002; Lau et al., 2009). For each condition, the M400 component was separately calculated by averaging over the individual trials from 250 ms to 600 ms following target word onset. The data were low-pass filtered at 10 Hz. All ERFs were baseline corrected based on a time window of −300 ms – 0 ms relative to target word onset. To facilitate comparison across participants the ERFs were transformed to a combine synthetic planar gradient representation (Bastiaansen & Knosche, 2000).

### Time-frequency analysis

Time-frequency analysis was first done for the time-window of −800ms to 0ms relative to the sentence’s target word onset, including only the blank delay period. Additionally, the M400 sensitive time-window after target word onset was considered for time-frequency analyses, including a window from 200 ms up to 700 ms. Power was estimated for each condition using fast Fourier transform for a frequency range of 8 Hz to 12 Hz for the alpha, and 16 Hz to 20 Hz for the beta frequency bands, with a Hanning-tapered 500 ms sliding window in time steps of 10ms.

### Source analysis

To estimate the sources of the oscillatory activity, the Dynamical Imaging of Coherent Sources (DICS) beamforming approach was applied to the data (Gross et al., 2001). The volume conduction model was constructed from the individual anatomical MRI as a single shell representation of the inside of the skull. This model was used to compute the forward model according to (Nolte, 2003). The initial co-registration between the headmodel and MEG sensors was achieved by manually identifying the anatomical landmarks of the nasion and two auricular fiducials, and was additionally refined, using the subject-specific three-dimensional digitised representation of the scalp, as obtained by a Polhemus digitizer. The source space was discretized into a three dimensional grid with a 6mm resolution. Source reconstruction was performed using a spatial filter, which was computed by combining the cross-spectral density (CSD) matrices from all three conditions (HC, MC, LC). The CSDs were computed using the Fast Fourier transform of the data with multitapering, with a center frequency of 10 Hz or 18 Hz for the alpha (averaged over the time window 540-0 ms, relative to target word onset) and beta (averaged over 450-0 ms) frequency band respectively. All visualizations are based on interpolated data onto the MNI template. The different conditions of the source reconstructed data were compared based on cluster-based permutation statistics as described below.

### Cluster-Based permutation statistics

Statistical evaluation was done using non-parametric cluster-based permutation tests (Maris & Oostenveld, 2007). First, we computed F-statistics to quantify the effect of context constraints (three levels: HC, MC, LC) for each sensor and time point. These F-statistics were used to define the clusters for the non-parametric statistical testing: clusters consisted of samples whose F-values were above threshold (threshold: F-value associated with a p-value of 5%) and were adjacent in space and time. Cluster-level statistics were computed by taking the sum of F-values within each cluster. The distribution of the cluster-level statistics under the null hypothesis was obtained by repeating this procedure for 5000 permutations of random relabeling of the conditions. Clusters whose test-statistics fell in the highest 5th percentile of its reference distribution were considered significant.

## RESULTS

### Behavioral performance

In order to confirm the participant’s attention to the experimental task, their performance was measured during catch trials that occurred after presentation of 20% of the sentences. The overall accuracy measures show a mean ceiling performance of 95% (SE = 4.65), 96% (SE = 4.42) and 97% (SE = 2.83) for the HC, MC and LC sentences respectively. There were no significant differences in accuracy with respect to the different conditions (Accuracy: F(2, 31) = 0.474, p = 0.627). This indicates that the participants were paying attention to the content of the presented sentences.

### Event-Related fields after target word onset

In this experiment, participants read words preceded by a context with different degrees of constraint (three context-constraint conditions: high, medium and low context constraints) while brain signals were recorded online. We first analyzed the effect of context constraints on the event-related fields upon target word presentation. Based on previous literature (DeLong et al., 2005; Kutas & Federmeier, 2011) we expected a monotonous relationship between cloze probability and the M400 component. Consistently, as can be seen from the amplitude fluctuations of the event-related activity (fig. 2), amplitude differences between the three conditions emerged within the typical M400 time-window. The M400 amplitude strength decreased with increasing cloze probability, such that the HC condition displayed the lowest M400 amplitude, followed by the MC and LC conditions. The cluster-based statistics revealed a main effect of context constraints on the M400 amplitude strength in a pre-defined time window of 250 ms to 600 ms after target word onset; this effect was most pronounced over a left lateralized cluster of sensors (fig. 2, cluster *p* = .002). The post-hoc contrasts (based on pairwise T-tests) revealed that the effect was mainly driven by a difference between HC vs. LC (*p* < .001) and HC vs. MC (*p* < .001). Although the M400 amplitude was smaller for MC than for LC condition, this difference was not significant (p = .057). These effects were overall in line with the current literature showing that the M400 amplitude reflects semantic retrieval and unification of the target word with the preceding context.

**Figure 2.**
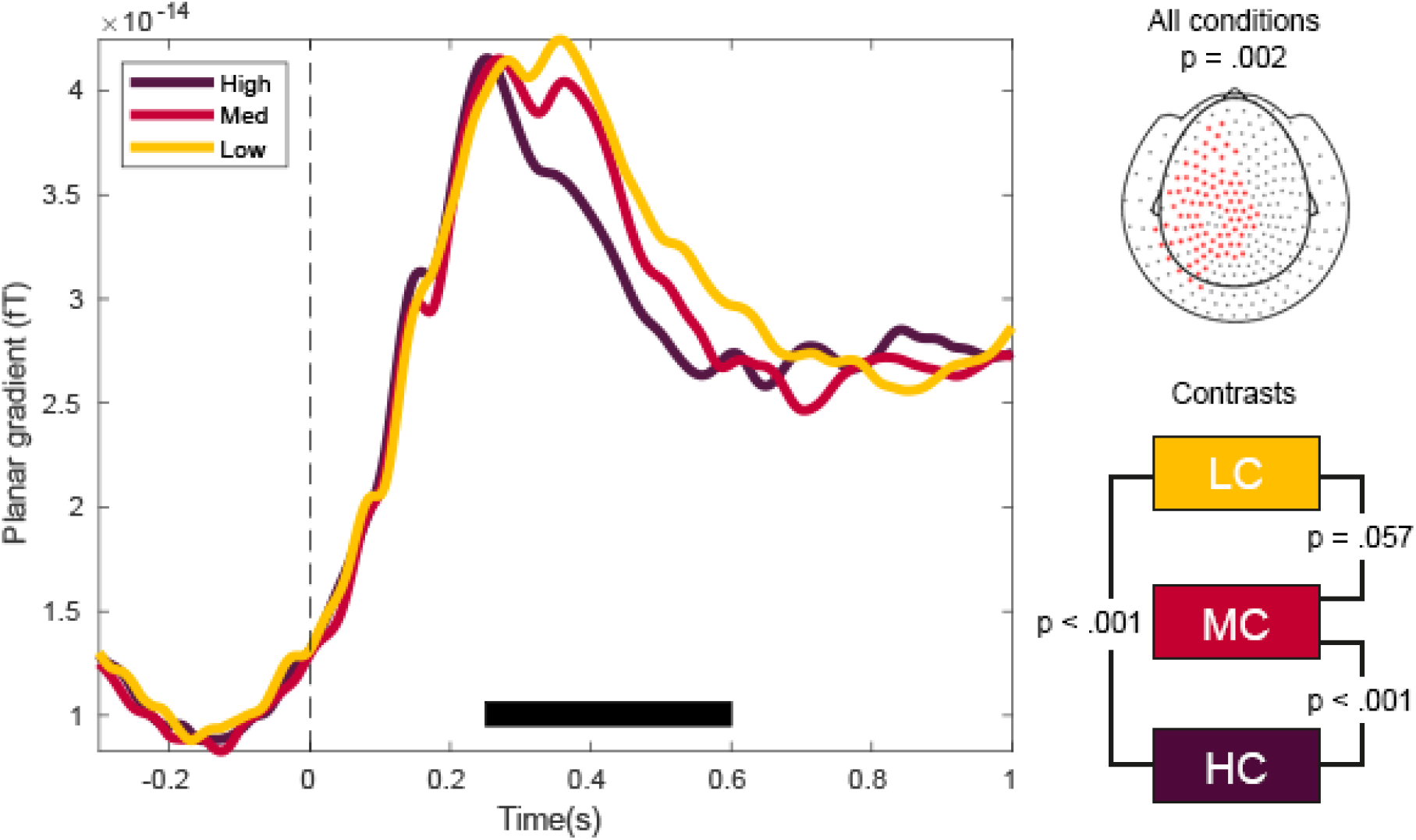
The event related fields of the M400 component after onset of a sentence’s target word. The striped line marks target word onset, the black bar indicates the significant time window. **Left**) The M400 amplitude in a time window of 250 ms to 600 ms is gradually modulated by the degree of context constraints, resulting in the lowest amplitude in HC, followed by MC and LC. **Right**) The upper topography is not only a display of the MEG sensors with respect to their position on the head, but also includes the helmet shape. The effect is most pronounced over left lateralized sensors.

### Alpha/Beta power modulations before target word onset

Next, we investigated the effect of content constraints on alpha and beta power modulations before target word onset. Based on earlier research (Wang et al., 2017), an effect was suggested to occur during the delay period, just before the display of the target word. The cluster-based statistics for the alpha (8-12Hz) frequency band revealed a significant difference between all three conditions with regard to power (*p* = .019) for a time-window of −540 ms to 0 ms relative to target word onset. This effect was most pronounced over a widespread set of sensors, including anterior, central and posterior sensors (fig. 3). Over these sensors, alpha power showed the strongest decrease for the MC condition, followed by HC and LC (fig. 3). The post-hoc contrasts of these conditions indicate that the power decrease is significantly different between HC vs. MC (*p* = .023), MC vs. LC (*p* < .001), and HC vs. LC. (*p* = .026). We performed source reconstruction to allow for a more detailed description of the brain areas involved in the observed sensor-level effect. Source-level statistical evaluation indicated that this effect was most pronounced in parietal areas, with a bias to the right hemisphere (fig. 4, p = .003, cluster-based corrected).

**Figure 3.**
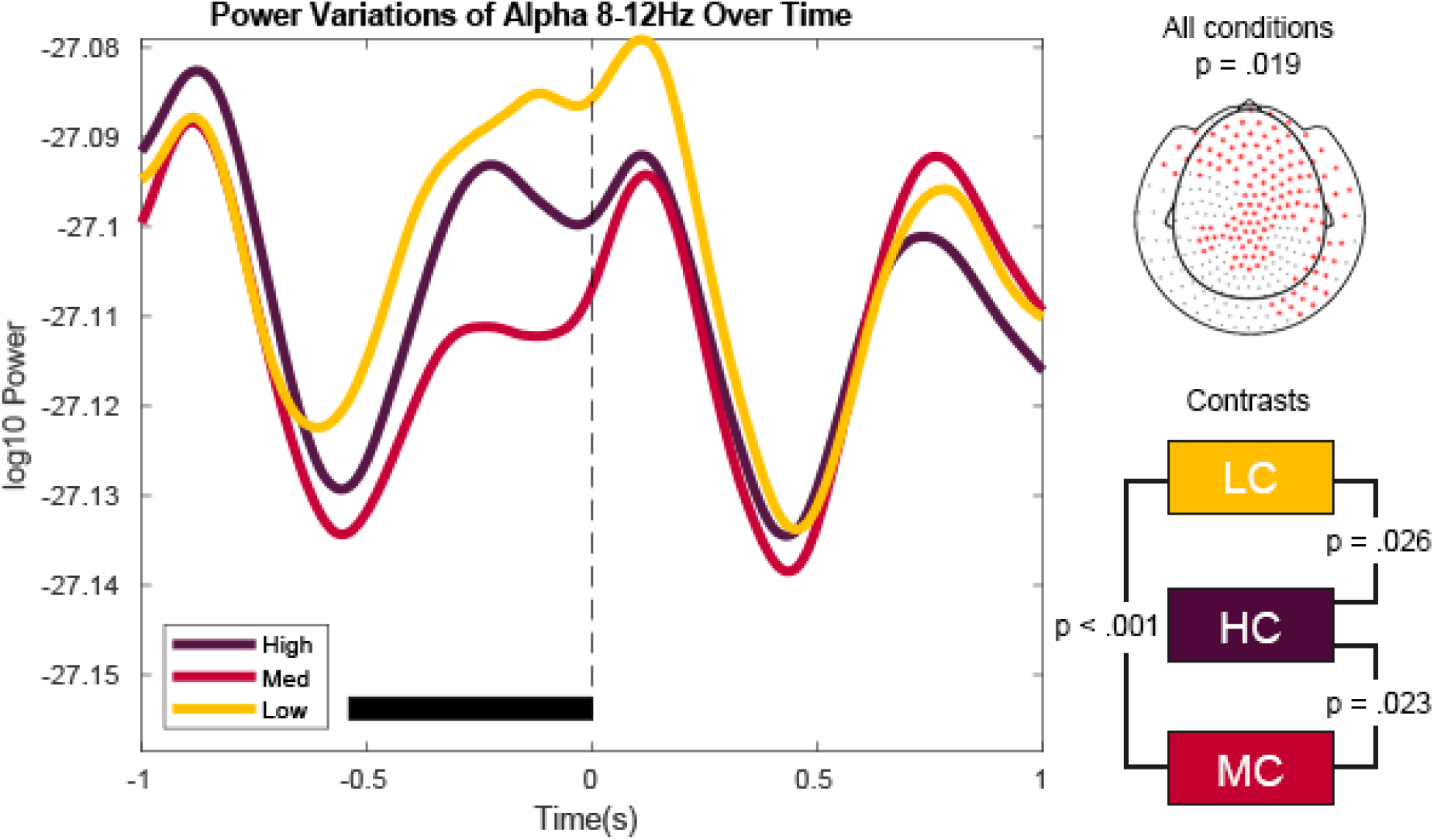
**Left)** Alpha power fluctuations over sensors where the effect of context constraint condition turned out to be significant. The striped line marks target word onset, the black bar indicates the significant time window. **Right)** Cluster results as obtained from the permutation statistics. The conditions differ significantly from each other and the effect is most pronounced over frontal and posterior sensors. The contrasts reveal significant differences only for HC vs. MC and MC vs. LC.

**Figure 4.**
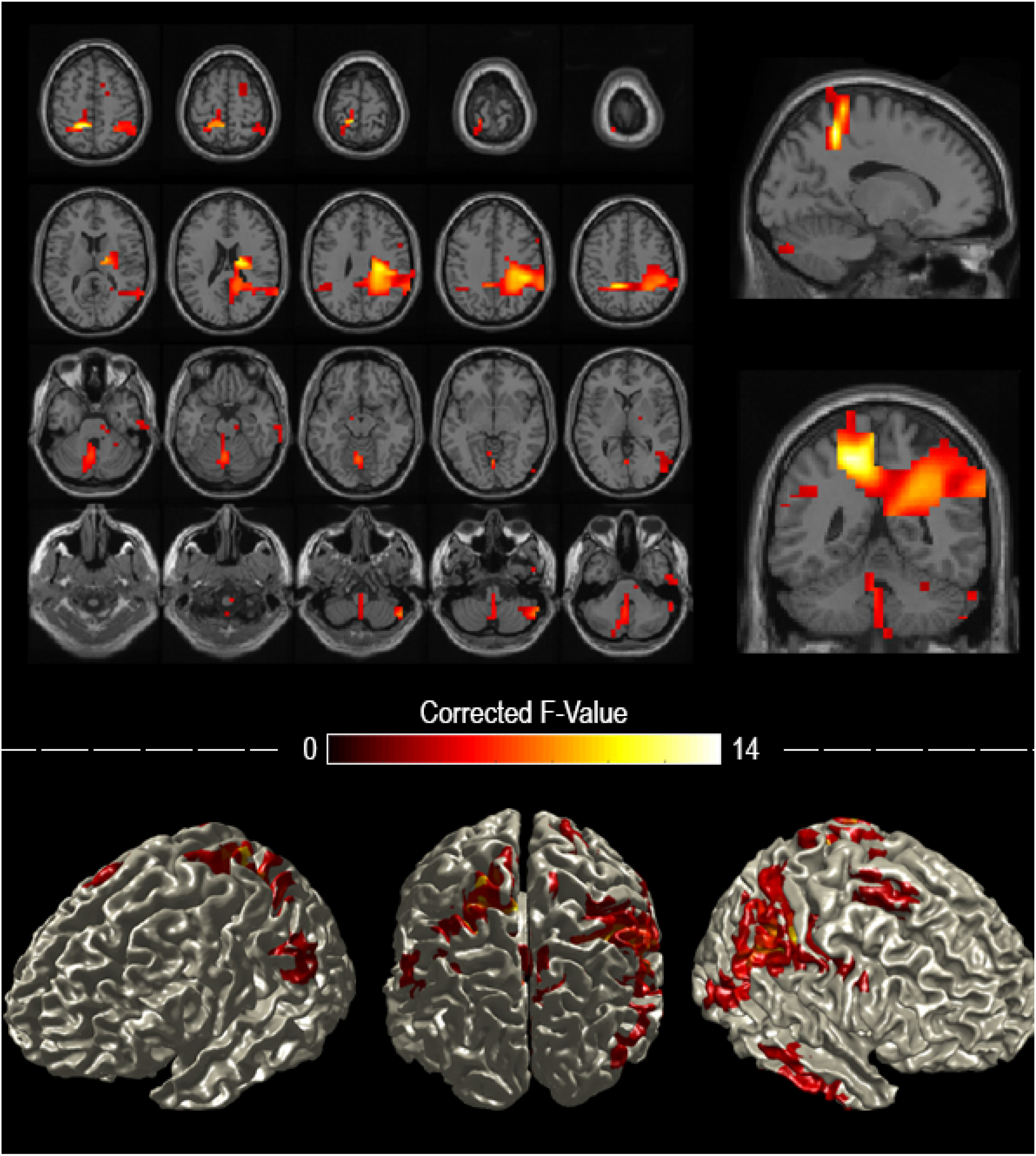
**Upper)** Reconstruction of the source statistics displayed as **(Left)** horizontal and **(Right)** sagittal as well as coronal slices for the alpha frequency band. **Lower)** Surface representation of the source statistics. The source statistics reveal that the effect of the context constraint manipulation is most pronounced over left and right parietal regions.

The sensor-level analyses in the low beta (16-20Hz) frequency band revealed a similar tendency as for the alpha results, though the effects were not significant (cluster with lowest p-value in cluster-based corrected statistics: *p* = .077, see supp. fig. 1). Source statistics in turn indicated a significant F-contrast across all conditions with the effect being most pronounced over a set of frontal and parietal areas, biased to left frontal cortex (supp. fig. 2, p = .002, cluster-based corrected). The power fluctuations were, similar to the results on alpha power, non-monotonically related to each other. The MC condition was again displaying the strongest decrease followed by HC and then the LC condition (supp. fig. 1). The overall results of the alpha and beta power decrease showed a non-monotonous behavior that cannot be easily reconciled with a prediction framework.

### Alpha/Beta power modulations after target word onset

To cover any oscillatory effects within the M400 sensitive time-window, alpha and beta power modulations were investigated as a function of context constraints after target word onset. The individual cluster-based statistics for both frequency bands revealed no significant difference between the three conditions with regard to power, within this particular time-window.

## DISCUSSION

The current study investigated the role of alpha and beta oscillations as a neural marker for sentence context constraints. Our results confirm the sensitivity of alpha and beta power to different levels of constraint, but we report the strongest decrease in alpha and beta power for the medium, followed by the high and then low context constraining condition. This effect was most pronounced over parietal sources in overlap for both frequency bands. In line with earlier findings, the M/N400 amplitude was monotonically modulated by the degree of constraint, resulting in the lowest amplitude for high, followed by medium and low context constraints. The results suggest that alpha/beta oscillations and the M/N400 component are neural markers that relate to distinct processes during sentence context evaluation.

In agreement with classic findings, our results show the M/N400 magnitude monotonically decreases with increasing context constraints, and this finding can be taken as support for M/N400 integration and/or predictability accounts (Kutas & Federmeier, 2011). In contrast, the non-monotonous alpha/beta power fluctuations do not support the predictability account. Our findings partially replicate previous results, showing that high context constraints induced a stronger alpha/beta power decrease than low constraint (Lewis et al., 2015; Rommers et al., 2017; Wang et al., 2017). Yet, the stronger alpha/beta power decrease for medium context constraints cannot be reconciled with previous interpretations of alpha/beta power in light of prediction. In addition to the differences in amplitude fluctuations between alpha/beta and the M400, our effects exhibit distinct topographical properties. The sources of the M400 have previously been localized to temporal as well as prefrontal areas, with a stronger prominence in the left hemisphere (see Lau et al., 2008). The topography of the M400 results in the present study is in line with these common findings. In contrast, the sources of the alpha/beta power changes cover largely parietal areas. The functional and topographical differences between these two neural markers therefore suggest two distinct underlying processes.

As an alternative to prediction related processes, alpha/beta oscillations could reflect other processes such as attention or working memory operations (Piai et al., 2015). The interaction between language processing and attention or memory operations is a continuously interactive process with information exchange that happens dynamically. Semantic context guides memory retrieval processes and constraints the set of target candidates that need to be retrieved and maintained (Levelt, 1989). These interactive operations in turn are potentially indicated by the alpha power modulations, which would not necessarily lead to the assumptions of a monotonous relationship between alpha/beta power and cloze probability. Indeed, alpha/beta oscillations have been previously related to the attentional gating and maintenance in working memory of task relevant items (Hanslmayr et al., 2012; Röhm et al., 2001; Roux & Uhlhaas, 2014). During sentence processing, alpha/beta oscillations could be then involved in the preselection and maintenance of lexical candidates (Piai et al. 2015; see also Piai et al., 2014). Based on our findings, we speculate that, depending on the degree of constraint, the landscape of priors generated by a language system changes and the interaction between maintenance supporting systems and the language system is modulated as well. The amount of possible lexical items to be maintained might differ, which would lead to distinct alpha/beta power modulations before target word onset. Compared to low or high context constraining sentences, intermediate context constraints may generate the highest competition between lexical candidates. In the high context constraining conditions, few items are competing, which would require low working memory demands. In low context constraining conditions, the sentential context is broad enough that the number of alternative candidates for sentence ending would be much higher than working memory capacities. Because of the broadness of possible lexical candidates, it is not possible to pre-activating only a limited number of linguistic items based on sentential context. We therefore speculate that working memory processes would not be engaged in this setting. Eventually, in medium context constraining settings, more distractors are to be maintained in working memory than in other context constrains conditions, which in turn is reflected by the stronger alpha power decrease.

In sum, using sentences with different context constraints that are matched on other linguistic variables like lexical frequency and word length, this MEG study shows that alpha/beta power in the course of the sentence is modulated by context constraints. However, the alpha/beta power decrease is strongest for medium constraining sentences, which defies previous interpretation of this marker in light of a prediction mechanism. The non-monotonic sensitivity of the alpha/beta power fluctuations to these different levels of constraints highlight the importance of including intermediate conditions in language research. While our results do not support a prediction framework account with regard to these oscillatory markers, a mechanistic account relating alpha/beta oscillations and the degree of sentence context constraint remains to be elaborated.

## Supplementary Materials

**Figure S1.**
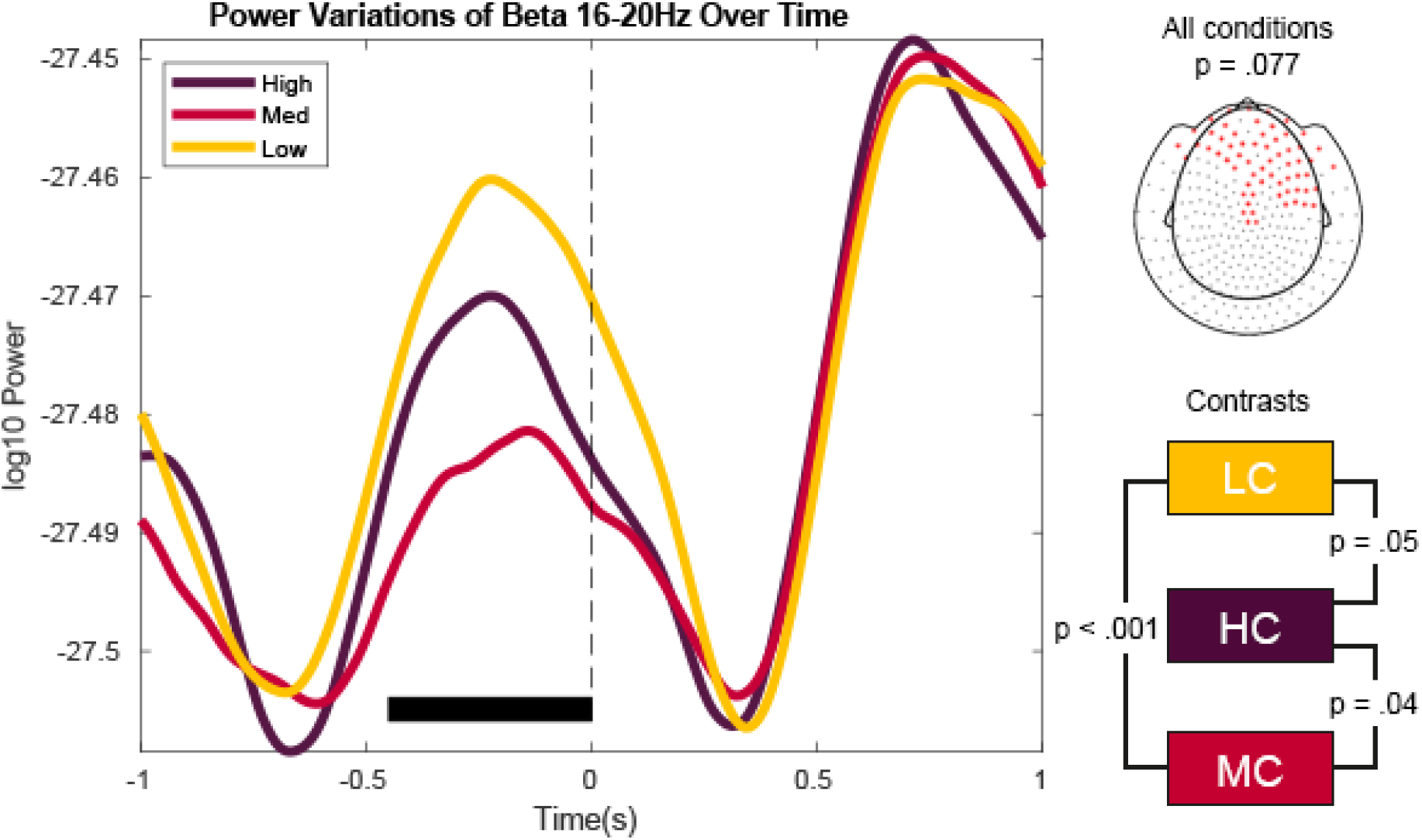
**Left)** Beta power fluctuations over sensors where the effect of context constraint condition turned out to be significant. The striped line marks target word onset, the black bar indicates the significant time window. **Right)** Cluster results as obtained from the permutation statistics. The conditions do not significantly differ from each other.

**Figure S2.**
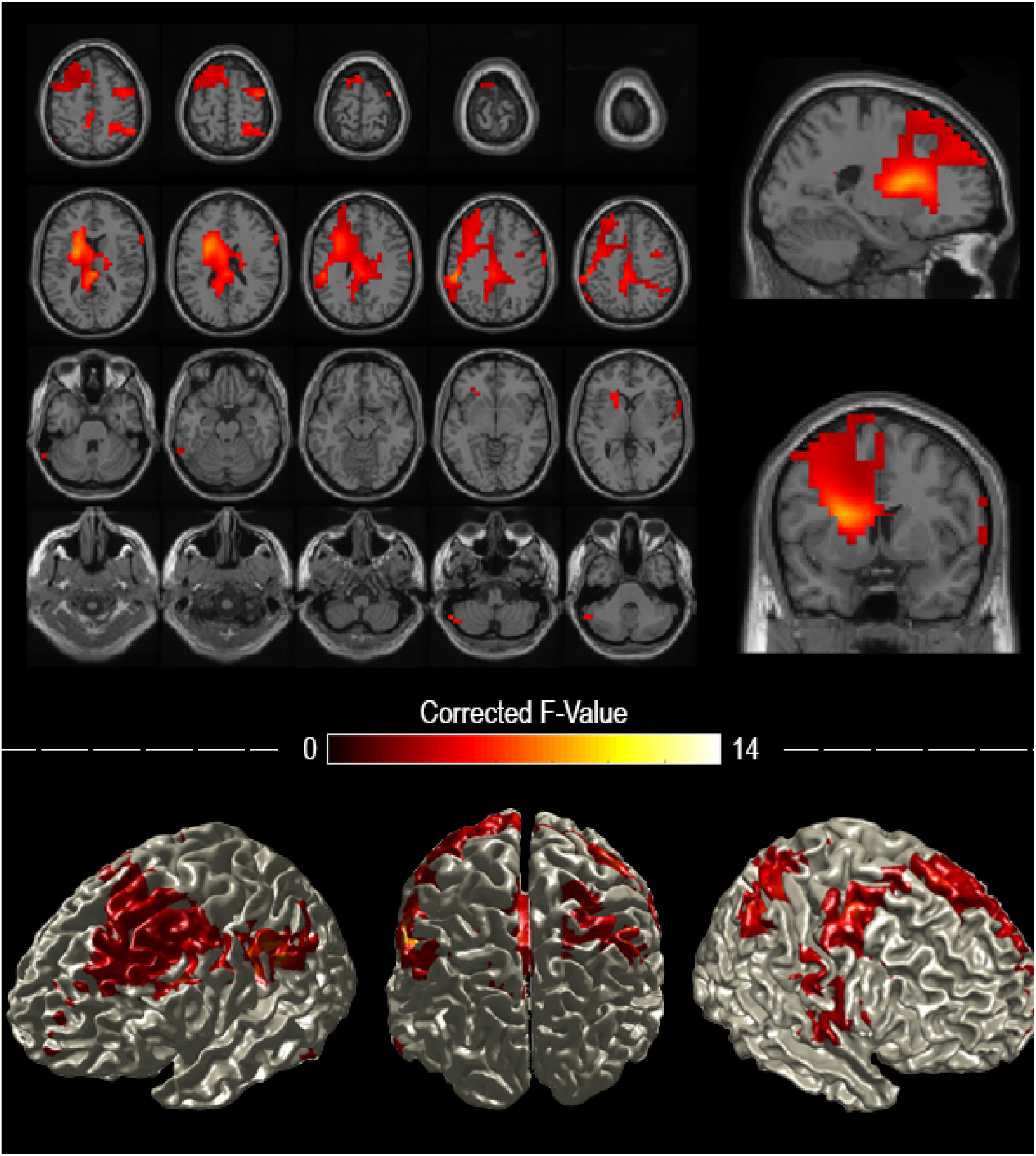
**Upper)** Reconstruction of the source statistics displayed as **(Left)** horizontal and **(Right)** sagittal as well as coronal slices for the beta frequency band. **Lower)** Surface representation of the source statistics. The source statistics reveal that the effect of the context constraint manipulation is most pronounced a distributed set of areas, but mainly involving left and right frontal areas and parietal regions.

